# The Delirium and Population Health Informatics Cohort study protocol: ascertaining the determinants and outcomes from delirium in a whole population

**DOI:** 10.1101/248559

**Authors:** Daniel Davis, Sarah Richardson, Joanne Hornby, Helen Bowden, Katrin Hoffmann, Maryse Weston-Clarke, Fenella Green, Nishi Chaturvedi, Alun Hughes, Diana Kuh, Elizabeth Sampson, Ruth Mizoguchi, Khai Lee Cheah, Melanie Romain, Abhi Sinha, Rodric Jenkin, Carol Brayne, Alasdair MacLullich

## Abstract

**Background:** Delirium affects 25% of older inpatients and is associated with long-term cognitive impairment and future dementia. However, no population studies have systematically ascertained cognitive function *before*, cognitive deficits *during*, and cognitive impairment *after* delirium. Therefore, there is a need to address the following question: does delirium, and its features (including severity, duration, and presumed aetiologies), predict long-term cognitive impairment, independent of cognitive impairment at baseline?

**Methods:** The Delirium and Population Health Informatics Cohort (DELPHIC) study is an observational population-based cohort study based in the London Borough of Camden. It is recruiting 2000 individuals aged ≥70 years and prospectively following them for two years, including daily ascertainment of all inpatient episodes for delirium. Daily inpatient assessments include the Memorial Delirium Assessment Scale, the Observational Scale for Level of Arousal, and the Hierarchical Assessment of Balance and Mobility. Data on delirium aetiology is also collected. The primary outcome is the change in the modified Telephone Interview for Cognitive Status at two years.

**Discussion:** DELPHIC is the first population sample to assess older persons before, during and after hospitalisation. The cumulative incidence of delirium in the general population aged ≥70 will be described. DELPHIC offers the opportunity to quantify the impact of delirium on cognitive and functional outcomes. Overall, DELPHIC will provide a real-time public health observatory whereby information from primary, secondary, intermediate and social care can be integrated to understand how acute illness is linked to health and social care outcomes.

## Background

Delirium is a severe neuropsychiatric syndrome mainly precipitated by acute illness, affecting at least 1 in 8 inpatients in industrialised countries.[1-3] Symptoms include acute onset of inattention, other cognitive deficits, altered level of consciousness, and psychosis.[4] Delirium has multiple adverse consequences, including higher mortality, longer hospital stay, and increased institutionalisation.[5-7] It is also highly distressing for patients, carers and staff.[8]

A range of studies have demonstrated that delirium is associated with future long-term cognitive impairment.[9-15] However, these have major methodological limitations, either:

(i) Delirium outcomes have been measured without pre-morbid baseline cognitive assessments, i.e. observed cognitive impairment at follow-up is confounded by undiagnosed pre-existing cognitive impairment (*Figure 1, top panel*);[13, 16] or
(ii) Delirium has been retrospectively ascertained, so detailed information on the features any delirium is lacking (*Figure 1, middle panel*).[9-12, 14, 15]

**Figure 1:**
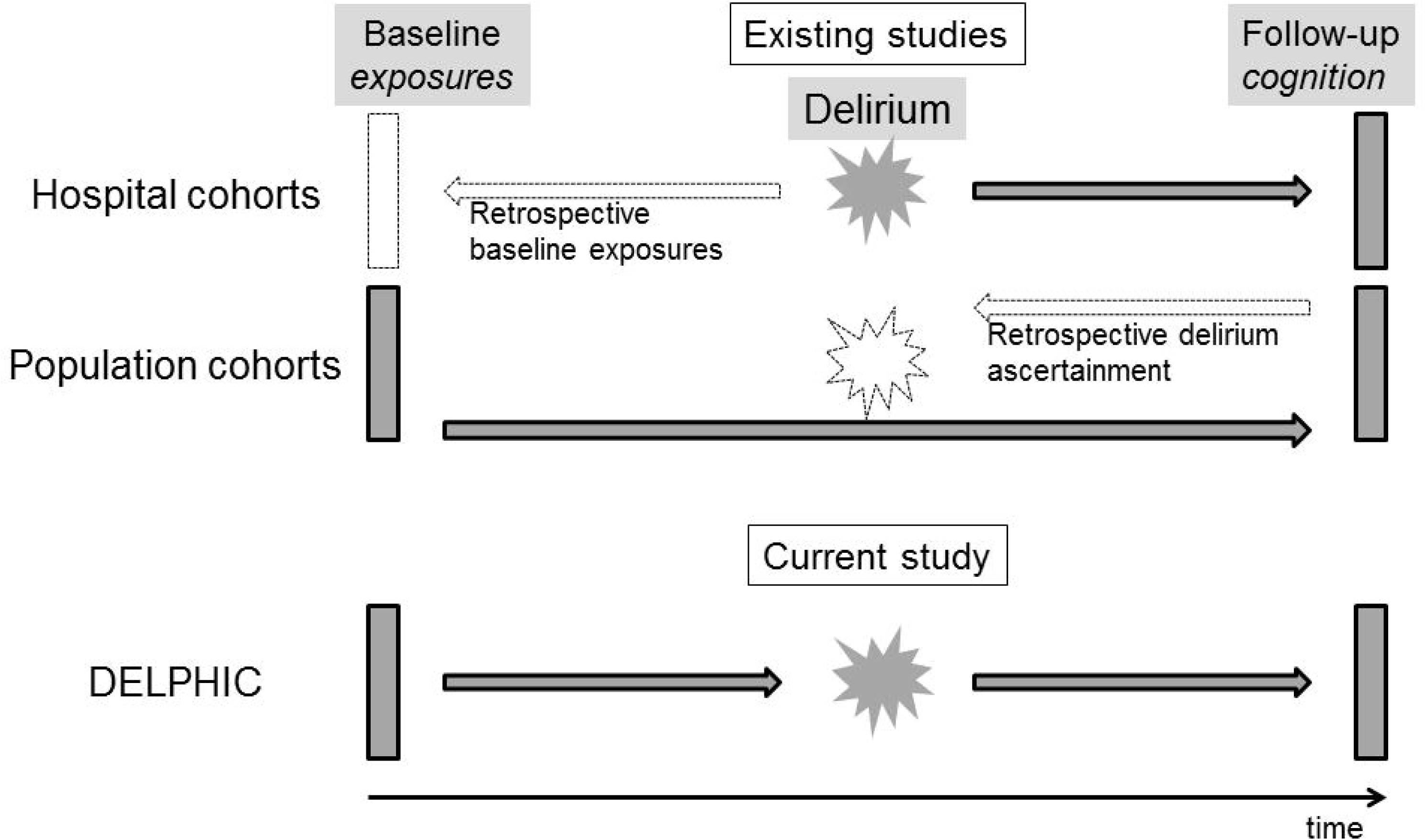
Studies examining delirium in relation to cognitive decline. *Top panel:* Hospitalised cohorts lack prospective measures of pre-morbid cognition. *Middle panel:* Population cohorts characterise cognition in community, retrospectively ascertaining delirium. *Lower panel:* A cohort prospectively tracking cognition before, during and after acute illness.

Thus, a critical gap is that no study has involved all three of the essential elements of (a) determining baseline cognitive function, then (b) ascertaining delirium prospectively, and then (c) assessing delirium’s impact on future long-term cognitive impairment. This is the key approach of the current study (*Figure 1, bottom panel)*.

Prospective ascertainment of delirium is important for several reasons. First, it is less subject to recall bias, which is common in the context of residual cognitive impairment. Second, prospective ascertainment allows for detailed assessment of the features of delirium. This is crucial because there are wide variations in the features of delirium, including severity, duration, and aetiology.[17] Such variations likely influence the risk of long-term cognitive impairment because delirium features affect other outcomes.[18-21] Finally, a focus on delirium would make possible the differentiation between its specific impact on cognitive outcomes, as opposed to the cognitive decline described in association with acute hospitalisation in general.[22-24]

To advance our understanding of the relationship between delirium and long-term cognitive impairment, we need to address the following question: does delirium, and its features (including severity, duration, and presumed aetiologies), predict long-term cognitive impairment, independent of cognitive impairment at baseline?

A definitive understanding of the natural history of delirium on risk of long-term cognitive impairment would have implications for identification and follow-up of patients at high risk of dementia, targeting acute treatment strategies, directing further research on mechanisms, and providing prognostic information to patients and carers.

## Methods

### Aim

To determine the impact of delirium, and its features, on the risk of long-term cognitive impairment in a population sample.

#### Objective 1

Recruit a population sample, the Delirium and Population Health Informatics Cohort (DELPHIC) (n=2,000).

#### Objective 2

Undertake a minimum of two community-based cognitive assessments, at baseline and two years.

#### Objective 3

Ascertain cumulative incidence of delirium (across community and hospital settings).

#### Objective 4

Quantify impact of delirium on change in long-term cognitive function.

#### Hypothesis

Incident delirium is associated with changes in global cognitive scores (pre-delirium compared to scores at two-year follow-up).

### Design

This is a prospective study of delirium and its features in relation to long-term cognitive impairment, recruiting a population sample and assessing cognition before, during and after delirium. Although DELPHIC is the scientific name for this study, recruitment will be known locally under the name: Long-term Information and Knowledge for Ageing (LINKAGE) Camden (www.linkage-camden.com).

### Population setting and sample

The sampling frame is geographically defined by the London Borough of Camden. Camden has 230,000 residents, 16,500 (7%) of whom are aged ≥70. It is one of the most socioeconomically varied areas of Europe. There is wide ethnic diversity; 16% of the population age ≥65 are non-White British according to the 2011 census.

All health care (primary and secondary) is commissioned by a single Clinical Commissioning Group (CCG) comprising 39 GP surgeries. The CCG is also co-terminus with provision of community rehabilitation (district nurses, physiotherapy, occupational therapy) and all community mental health services are provided by Camden and Islington NHS Foundation Trust. Social services and public health are provided by the local authority directly. Camden is served by two acute hospitals, University College Hospital (UCH) and the Royal Free Hospital (RFH). These are the main sites for clinical ascertainment of delirium. Together, this represents the opportunity to determine and integrate the entire health and social care usage of participants over the follow-up period.

### Eligibility

#### Inclusion criteria

Resident in Camden, registered with a Camden GP, age ≥70 years.

#### Exclusion criteria

Severe hearing impairment or aphasia, unable to speak English sufficiently to undertake any cognitive assessment, terminal phase of illness.

#### Participant characteristics

Participants are recruited from the 39 general practices in sequence, initially targeting the larger practices and purposively including a wide socio-economic distribution. Two care homes are also involved, with the aim of including a representative proportion of the Camden population in residential and nursing care (approximately 5% of sample). Patients previously under care of University College and Royal Free Hospitals are also approached. Patients known to Camden Memory Service are invited (Figure 2).

**Figure 2:**
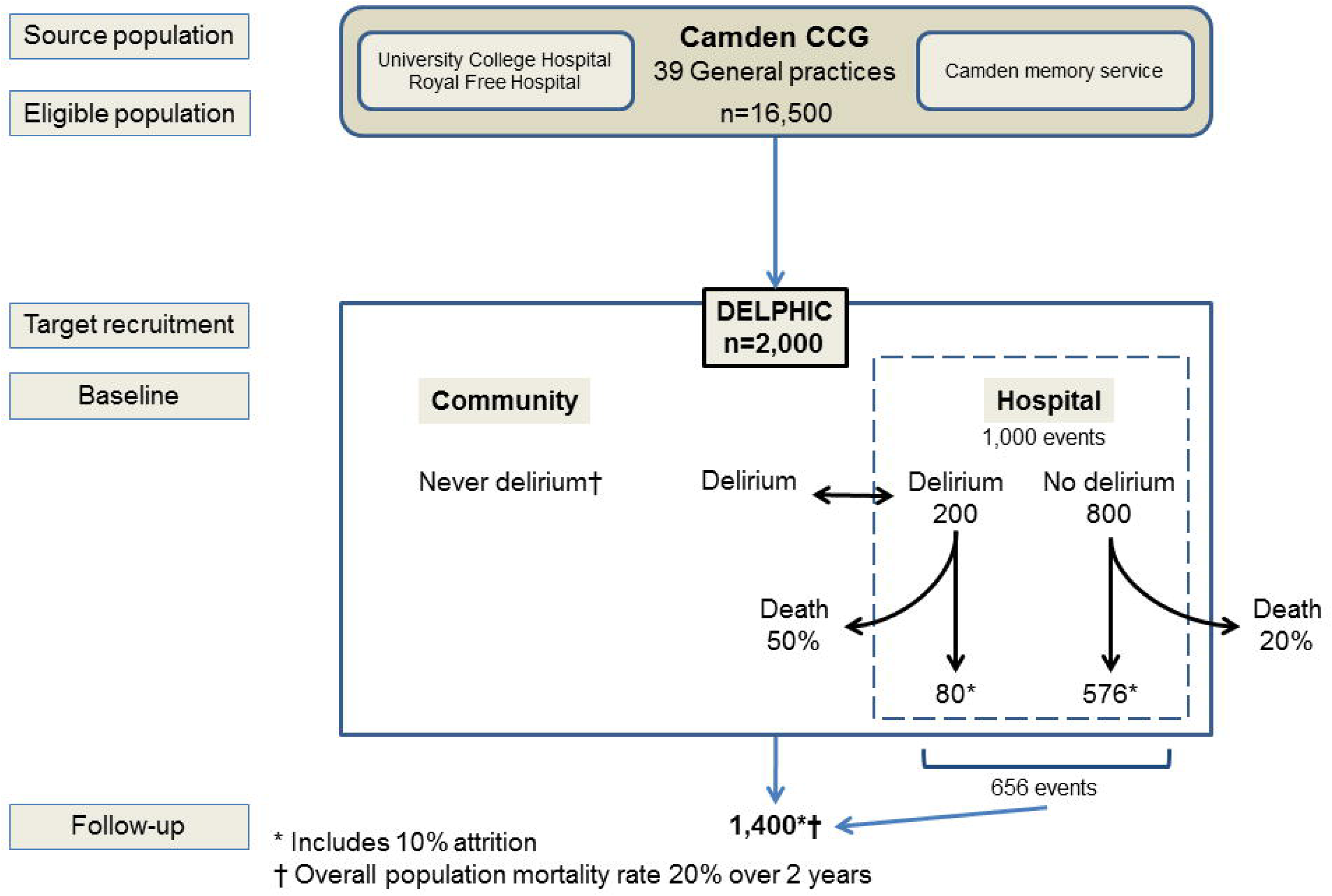
Study flow diagram showing recruitment sources, follow-up and expected attrition over two years.

Through approaching participants from GP registers (generally healthy), those known to hospital services (enriched for comorbidity) and memory clinics (enriched for dementia) in a 8:1:1 ratio, the target sample is expected to match the age structure of the 2011 census, as well as have the expected distribution of cognitive function in the population using data from the Cognitive Function and Ageing Study II.[25]

### Consent

Consent is obtained in line with the principles of Good Clinical Practice. At the start of the study, participants will also be asked to consider their wish to remain in the study in the event that they lose capacity during the study period.

This study will involve participants who lack capacity because it is specifically about cognitive impairment and its acute and chronic determinants. At this project's heart is the recognition that older adults with cognitive impairment have high health and social care needs, but with little research that understands the impact of having cognitive impairment as individuals move between primary, secondary and intermediate care. This research is designed to investigate directly the needs of adults with cognitive impairment, including those unable to give consent for themselves. To not include such participants introduces bias into the research and leaves clinicians and policy makers with no research data to improve care for older patients unable to give consent and invalidate the study almost entirely.[26]

For those lacking capacity, a consultee will be sought. Consultees will be routinely sought for all participants, including those with capacity who give consent for this. Where necessary, this may include the GP acting as a professional consultee in line with Section 32 of the Mental Capacity Act. In many cases, consultees will provide important collateral information and their continued involvement will be encouraged. Capacity can fluctuate during delirium and dementia. Where an event occurs that is part of the study (e.g. hospitalisation), consultees will be sought if the participant is unable to give continued consent as appropriate. Thereby, it is intended that for those individuals who lose capacity at any stage the research will continue to be able to participate under the terms of the Mental Capacity Act 2005.

### Data to be collected

All participants undergo a baseline assessment, repeated two years later. Data collection started in March 2017 and end of the follow-up period will be in March 2020. Those admitted to hospital will be seen throughout their admission, usually daily. Participants discharged from hospital, and those deemed to be at high risk for delirium will be contacted every two months by telephone in order to estimate incidence of delirium in the community using the Informant Assessment of Geriatric Delirium[27] and quantify trajectories of recovery after delirium (Figure 3). The data acquired in each setting are summarised in Table 1.

**Figure 3:**
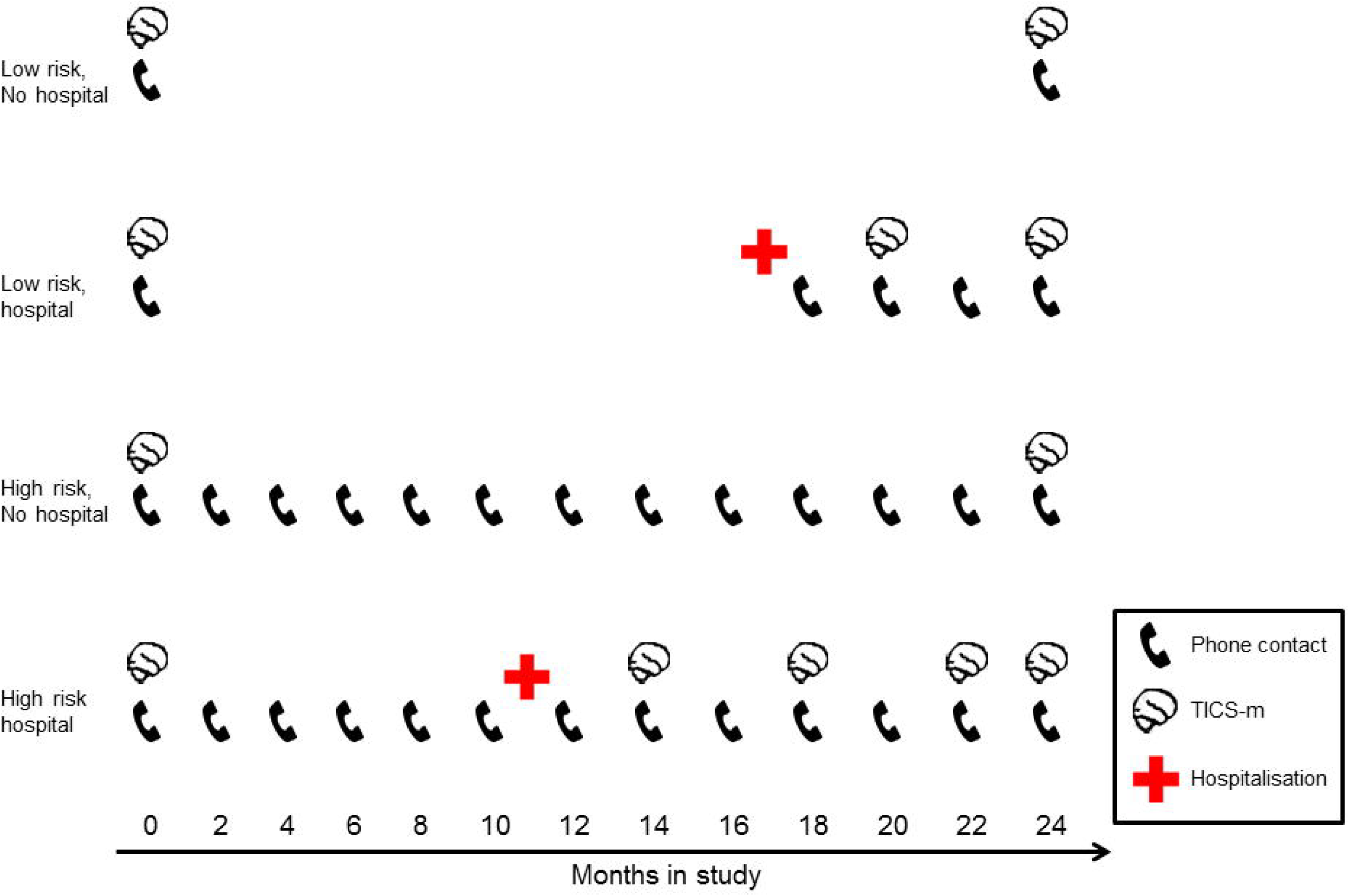
Schematic showing telephone contacts and cognitive testing in four examples, depending on baseline risk for delirium. Both the number of contacts and cognitive assessments increase in the event of hospitalisation.

**Table 1.**
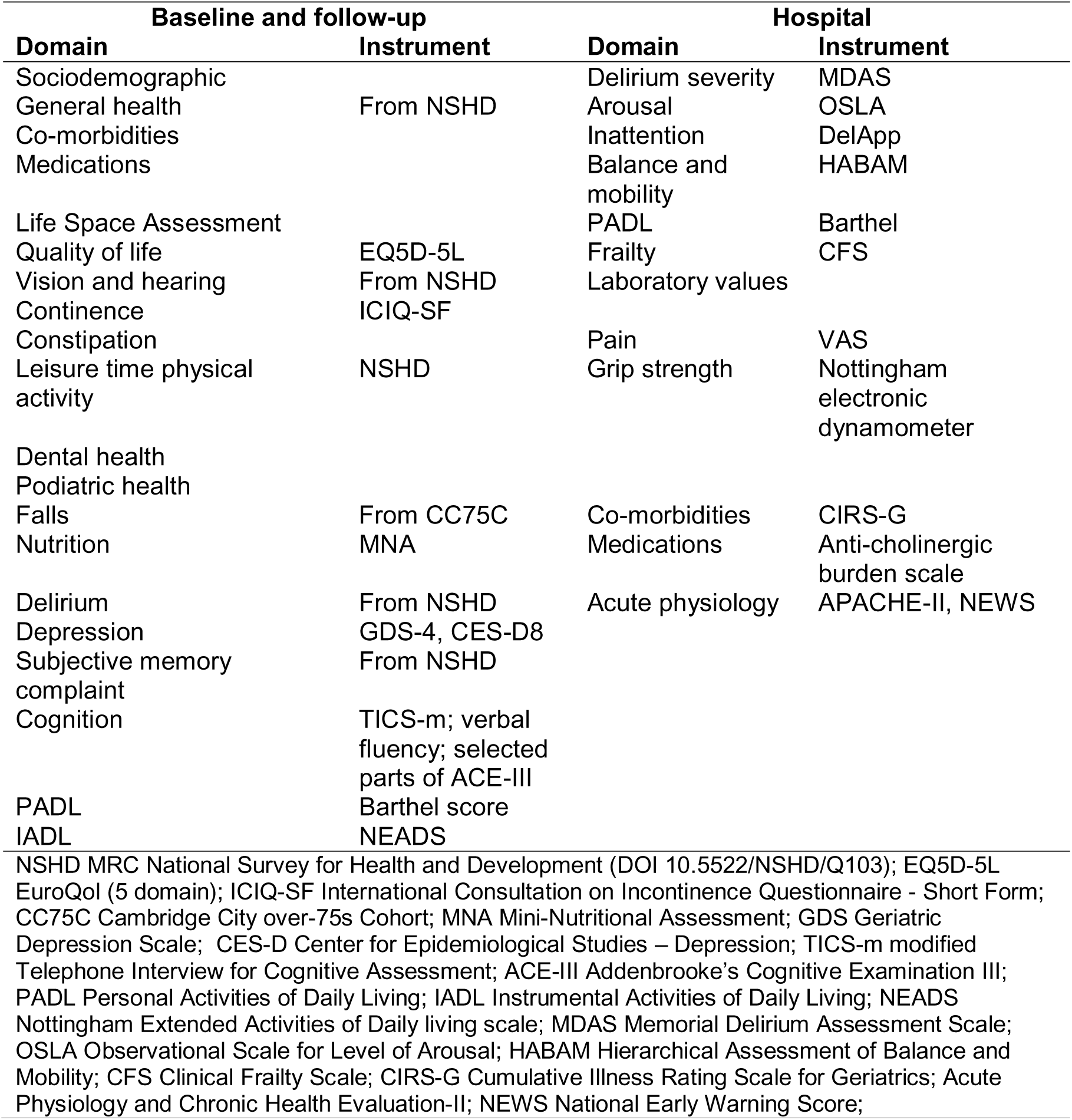
Summary of assessments

### Community assessments

These are mainly undertaken by telephone, though some participants are seen at home or in the Bloomsbury Centre for Clinical Phenotyping. The whole sample is assessed at baseline and two years later (primary outcome measure). In addition, a 12.5% subsample (n=250) of the most cognitively impaired (highest-risk for delirium) is being proactively monitored by the research team (Figure 3).

The baseline contact comprises:

- Consent for involvement in DELPHIC, specifically including: hospital assessments in the event of acute illness (particularly if capacity is impaired through delirium or dementia at subsequent contacts); record linkage of electronic health data in primary and secondary care
- If capacity to consent is impaired, a consultee declaration will be sought, in line with NHS Health Research Authority guidance
- Nomination of consultees
- Administration of the Modified Telephone Interview for Cognitive Status (TICS-m),[28, 29] plus verbal fluency and other memory tests from the Addenbrooke’s Cognitive Examination III
- Other measures of health and wellbeing, including: general health, co-morbidities, medications, health behaviours, hearing, vision, quality of life, continence, falls, depression, personal and instrumental activities of daily living (Table 1)

TICS-m is a widely-used, validated test which takes 10 minutes to administer over the telephone or in person.[30] Cognitive domains measured include: orientation, concentration, delayed recall, language, praxis, calculation, verbal comprehension. It is scored out of 50 points and has a normal distribution in population samples of older persons.[31] While severe hearing impairment would preclude assessment with TICS-m, it is possible to test individuals with severe visual and/or motor impairments.

### Proactive telephone contact for highest risk

Age and baseline cognitive impairment are the strongest risk factors for delirium. A subsample of the 12.5% most cognitively impaired (n=250) (Table 2) are selected for enhanced delirium surveillance, before and after any hospitalisations (Figure 3). This strategy has previously been used for ascertaining other acute events in population samples, e.g. falls.[32, 33] This allows for the most complete understanding of how delirium develops and patterns of recovery across healthcare settings, thereby challenging the assumption that most delirium presents to acute hospitals.

**Table 2.**
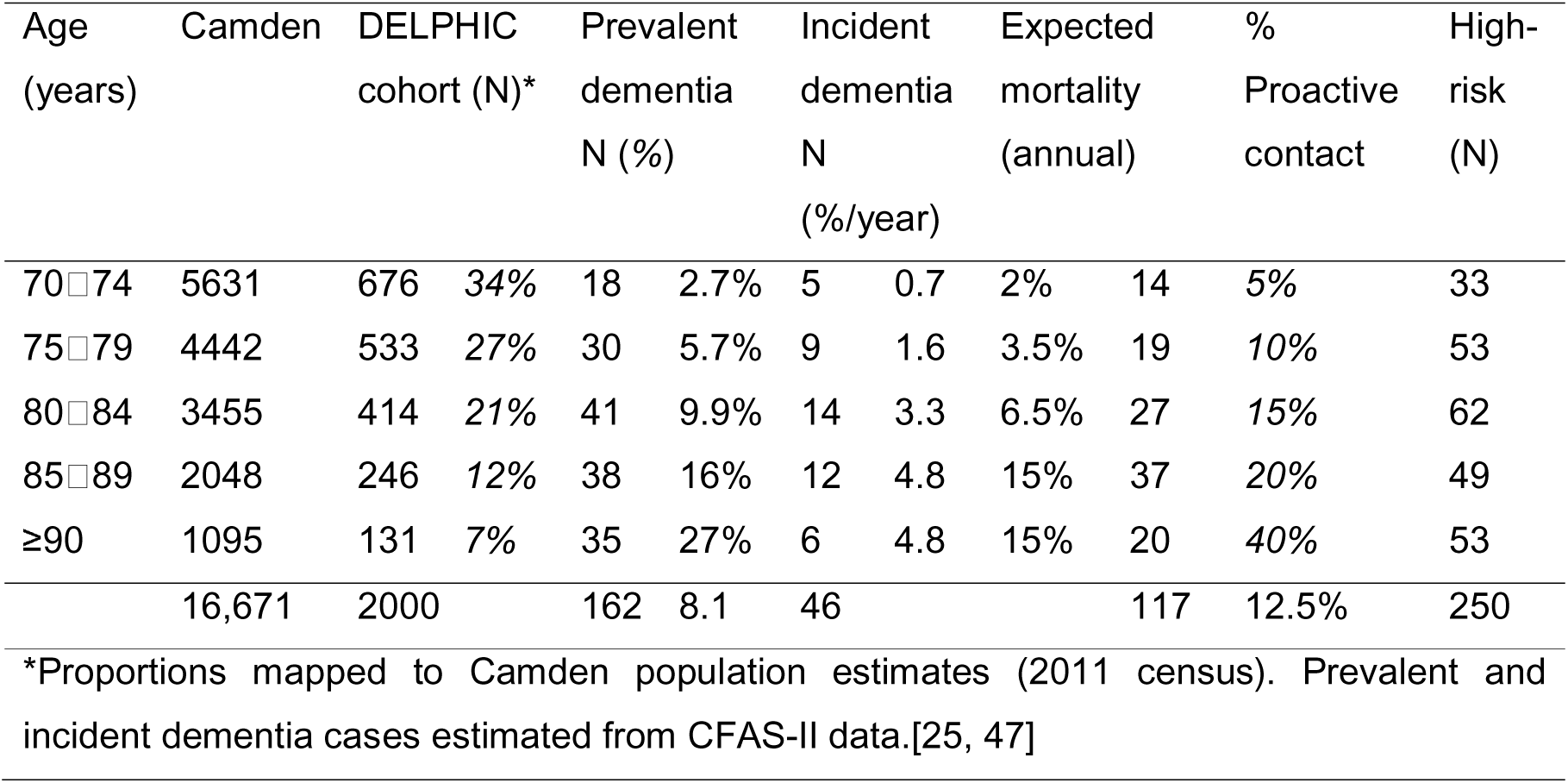
Age structure of DELPHIC in relation to dementia prevalence, hospital presentation rate and sample for proactive contact.

The research team undertake telephone contacts each day (Monday-Friday), covering the subsample of 250 participants and/or their nominated trusted advisors (consultees) every two months (Figure 3). Each contact has the following purposes:

- Assess any new health problem (every two months).
- Assess any delirium symptoms using the validated Informant Interview for Geriatric Delirium (I-AGeD[27]). This provides key information on community delirium both before hospitalisation and tracks recovery after hospital (every two months).
- Repeated TICS-m in participants after discharge (every four months) (Figure 3).

### Hospital assessments

Admissions lists are screened Monday-Friday, identifying participants who have been admitted (emergency and elective). Specific audit data from Camden practices, along with reports from NHS Digital, indicate that the admission rate in this age group is up to 20/1000/month. This amounts to 1,000 events over the two year study period, with the highest-risk being admitted recurrently.

Identified admissions are assessed for delirium. Data recorded includes information collected through usual clinical care:

- Demographic: age, sex, education, place of residence, co-resident support
- Clinical: admission details, physiological measurements (National Early Warning Scores (NEWS)), illness severity scores (Acute Physiology and Chronic Health Evaluation (APACHE) II (minus arterial blood gas)), medications.
- Delirium: general cognition (TICS-m), MDAS, arousal, attention (including the DelApp[34]), functional balance and mobility, aetiological factors.

Participants admitted to UCH or RFH are seen every weekday. Relevant clinical data from out-of-hours (including weekends and participants discharged before assessment) maximise ascertainment using validated method for detecting delirium from medical notes and interviews with ward staff and family.[35] Hospitalised participants will be followed up at St Pancras Hospital if they are discharged to bed-based rehabilitation. Participants at St Pancras will be seen a minimum of twice a week.

After hospitalisation, participants (and/or proxies) will continue to be proactively contacted as described above. There will be up to five additional occasions for administering TICS-m, adding longitudinal information on trajectories to recovery or persistent delirium (*Figure 3*).

Delirium ascertainment is supervised by DD, with difficult cases used for ongoing training and knowledge sharing. Complex cases are adjudicated on a monthly basis with input from specialist old age liaison psychiatry (ES). The final delirium variables (incidence, duration, severity, aetiology) will be derived by an expert consensus panel, blinded to outcome data. All available inpatient assessments, telephone contacts, electronic hospital and GP records collected for data linkage will be used.

### Additional data sources

Participants are asked to consent for access to their NHS medical and social care notes, including data from GP, community mental health, community rehabilitation and social services records throughout the duration of the study. The facility to do this comes from the Camden Integrated Digital Record.

### Statistical methods

#### Power calculations for hypothesis

Figure 2 shows the expected mortality for hospitalised persons (delirium and non-delirium).[5, 9, 23, 36] Overall mortality in the population aged ≥70 years is 12%/year (Office for National Statistics). Attrition from causes other than death is estimated as 10%. Calculations used the *sampsi* command in Stata (version 12.1), and assume α=0.05 and β=0.9. The most conservative estimate of TICS-m standard deviation reported in the literature (SD=7.2) was used.[31]

A clinically significant change in the two-year total TICS-m score would be 6 (out of 50) points for hospitalised delirium patients (change within-person for incident delirium cases) (hypothesis) and 3 points for hospitalised persons without delirium (compared to delirium cases).[23, 24] This effect size is consistent with other studies using this type of primary outcome where incident delirium was associated with change in global cognition scales of 2.5 (out of 28) points (Blessed Information-Memory-Concentration test)[11] and 4 (out of 30) points (Mini-Mental State Examination).[10]

The resulting sample size is n=215 to detect the primary outcome (within-person change in the incident delirium group) and n=431 to compare differences between hospitalised participants without delirium. This allows a margin to assess 70% of admissions or a 50% overestimate of delirium cases, and still be sufficiently powered.

### Statistical analyses

#### Outcome

TICS-m score at follow-up

#### Exposures

**Main exposure:** <u>delirium</u> (severity (MDAS scores); duration (days), modelled as a time-varying covariate across the whole study period); aetiology (four categories: infective/inflammatory; metabolic; pharmacological; other)

**Confounders:** <u>baseline</u>: TICS-m score at baseline, age, sex, ethnicity (three categories), education level (three categories); <u>illness severity</u>: APACHE II at admission and daily total NEWS scores

#### Analyses

- Delirium incidence will be expressed as an annual rate, and described stratified by age.
- Linear regression, where outcome is TICS-m score at two years, will be used (hypothesis). 14 parameters are proposed, this can be accommodated by a follow-up sample of 1,400.
- Timing of delirium is important and delirium variables will be analysed as time-varying covariates, where this can be considered ‘time at risk’ for change in TICS-m score. This also allows the effects of recurrent delirium to be assessed and is flexible for differences in time intervals between delirium occurrences and follow-up.
- More detailed analyses of trajectories in relation to repeated TICS-m scores will be possible using random-effects models.[37]
- Data missing at random will be treated using multiple imputation.
- Where appropriate, shared parameters models may jointly link random-effects models with survival analyses to account for attrition due to death.[38]

#### Patient and public involvement

Patient and public involvement operates through the formation of a PPI group with input throughout the course of the study. The group is drawn from interested persons in Camden, including those involved with the CCG, Age UK, Carers UK, Alzheimer’s Society. The PPI group is involved in refining the study documentation (PIS, consent forms), recruitment strategies as well as dissemination of findings. The group meets every four months, where new study questions and modalities of data collection are considered.

### Discussion

The DELPHIC study represents an opportunity to characterise prospectively the impact of delirium on long-term cognitive impairment. It will provide a definitive estimate of cumulative incidence of delirium across settings in a whole population. Prospectively linking a community sample with hospitalisations will lead to new knowledge on pathways to longterm cognitive impairment, overcoming the limitations of previous studies in selected samples. DELPHIC also offers an opportunity to explore mechanisms by providing a population framework to nest representative samples testing hypotheses from experimental studies.[39, 40]

With respect to other cohort studies, DELPHIC is closely related to CFAS-DECIDE, where the delirium ascertainment protocols were developed in conjunction.[41] The community assessments have overlap with measures undertaken in the MRC National Survey for Health and Development.[42, 43] In ascertaining both delirium and dementia, DELPHIC will be a contributing cohort to the Dementias Platform UK.

DELPHIC will lead to a resource for insights into the delirium-dementia relationship from its biological underpinnings through to the public health implications. A systematic characterisation of temporal patterns of acute illness, hospitalisation, delirium and cognitive outcomes is urgently required.[44] DELPHIC will also inform how underlying dementia influences the incidence and detection of delirium, by adding empirical data to the clinical uncertainties surrounding delirium superimposed on dementia.[45, 46] By analysing whole population transitions of cognitive function in older people across healthcare settings, DELPHIC will lead to greater understanding of progression of cognitive impairments in ageing.

## Abbreviations

APACHE II: Acute Physiological and Chronic Health Evaluation
DELPHIC: Delirium and Population Health Informatics Cohort
DSM-IV: Diagnostic and Statistical Manual, 4^th^ edition
I-AGeD: Informant Assessment for Geriatric Delirium
LINKAGE-Camden: Long-term Information and Knowledge for Ageing in Camden
MDAS: Memorial Delirium Assessment Scale
NEWS: National Early Warning Score
RFH: Royal Free Hospital
TICS-m: Telephone Interview for Cognitive Status - modified
UCH: University College Hospital

## Declarations

### Ethics approval and consent to participate

This study has received approval from the Camden and King’s Cross Research Ethics Committee (16/LO/1217) and the Health Research Authority. Written informed consent is obtained from participants. In those individuals found to be without capacity to give full written informed consent, a personal consultee is identified and their advice sought regarding participation.

## Consent for publication

Not applicable

## Availability of data and materials

Metadata on request, data sharing applications through the MRC Unit for Lifelong Health and Ageing at UCL.

## Competing interests

The authors declare that they have no competing interests Funding DELPHIC is funded by the Wellcome Trust through an Intermediate Clinical Fellowship and a Wellcome-Beit Prize to DD (WT107467).

## Authors' contributions

DD drafted the manuscript, acquired funding, JH, HB, KH, MWC, FG, RM, KLC, MR, AS, RJ helped to devise local procedures, NC, DK, CB, AM advised on the academic aspects as did SR who also drafted SOPs. All authors read and approved the final manuscript.

## Acknowledgements

We would like to acknowledge the help and support of all participating health and social care professionals in Camden. We give our continued thanks to all the DELPHIC/LINKAGE participants and their families for their time and involvement in our study.

### Key definitions

**Delirium:** DSM-IV

**Duration:** Days in delirium, determined by consensus of all data obtained: direct assessment, informant interview, hospital and community clinical records.

**Severity:** Serial Memorial Delirium Assessment Scale (MDAS) scores associated with each delirium episode.

**Aetiology:** Principal precipitating causes (infective/inflammatory; pharmacological; metabolic; other), determined by consensus of all data obtained.

**Cognitive function:** Modified Telephone Interview for Cognitive Status (TICS-m) score.

## References

1. Ryan DJ, O'Regan NA, Caoimh RO, Clare J, O'Connor M, Leonard M, McFarland J, Tighe S, O'Sullivan K, Trzepacz PT et al: Delirium in an adult acute hospital population: predictors, prevalence and detection. BMJ open 2013, 3(1):e001772.

2. Maclullich AM, Anand A, Davis DH, Jackson T, Barugh AJ, Hall RJ, Ferguson KJ, Meagher DJ, Cunningham C: New horizons in the pathogenesis, assessment and management of delirium. Age Ageing 2013, 42(6):667-674.

3. Jackson TA, Gladman JRF, Harwood RH, MacLullich AMJ, Sampson EL, Sheehan B, Davis DHJ: Challenges and opportunities in understanding dementia and delirium in the acute hospital. PLoS medicine 2017, 14(3):e1002247.

4. Meagher DJ, Moran M, Raju B, Gibbons D, Donnelly S, Saunders J, Trzepacz PT: Phenomenology of delirium: Assessment of 100 adult cases using standardised measures. British Journal of Psychiatry 2007, 190(FEB.):135-141.

5. Witlox J, Eurelings LS, de Jonghe JF, Kalisvaart KJ, Eikelenboom P, van Gool WA: Delirium in elderly patients and the risk of postdischarge mortality, institutionalization, and dementia: a meta-analysis. JAMA: the journal of the American Medical Association 2010, 304(4):443-451.

6. Fong TG, Jones RN, Marcantonio ER, Tommet D, Gross AL, Habtemariam D, Schmitt E, Yap L, Inouye SK: Adverse Outcomes After Hospitalization and Delirium in Persons With Alzheimer Disease. Annals of Internal Medicine 2012, 156(12):848-U121.

7. Dani M, Owen LH, Jackson TA, Rockwood K, Sampson EL, Davis D: Delirium, frailty and mortality: interactions in a prospective study of hospitalized older people. J Gerontol A Biol Sci Med Sci 2017.

8. Partridge JS, Martin FC, Harari D, Dhesi JK: The delirium experience: what is the effect on patients, relatives and staff and what can be done to modify this? International journal of geriatric psychiatry 2013, 28:804-812.

9. Davis DH, Barnes LE, Stephan BC, MacLullich AM, Meagher D, Copeland J, Matthews FE, Brayne C, Function MRCC, Ageing S: The descriptive epidemiology of delirium symptoms in a large population-based cohort study: results from the Medical Research Council Cognitive Function and Ageing Study (MRC CFAS). BMC Geriatr 2014, 14:87.

10. Davis DH, Muniz Terrera G, Keage H, Rahkonen T, Oinas M, Matthews FE, Cunningham C, Polvikoski T, Sulkava R, MacLullich AM et al: Delirium is a strong risk factor for dementia in the oldest-old: a population-based cohort study. Brain 2012, 135 (Pt 9):2809-2816.

11. Fong TG, Jones RN, Shi P, Marcantonio ER, Yap L, Rudolph JL, Yang FM, Kiely DK, Inouye SK: Delirium accelerates cognitive decline in Alzheimer disease. Neurology 2009, 72(18):1570-1575.

12. Gross AL, Jones RN, Habtemariam DA, Fong TG, Tommet D, Quach L, Schmitt E, Yap L, Inouye SK: Delirium and long-term cognitive trajectory among persons with dementia. Arch Intern Med 2012, 172(17):1324-1331.

13. Pandharipande PP, Girard TD, Jackson JC, Morandi A, Thompson JL, Pun BT, Brummel NE, Hughes CG, Vasilevskis EE, Shintani AK et al: Long-term cognitive impairment after critical illness. New England Journal of Medicine 2013, 369(14):1306-1316.

14. Davis DH, Muniz-Terrera G, Keage HA, Stephan BC, Fleming J, Ince PG, Matthews FE, Cunningham C, Ely EW, MacLullich AM et al: Association of Delirium With Cognitive Decline in Late Life: A Neuropathologic Study of 3 Population-Based Cohort Studies. JAMA Psychiatry 2017, 74(3):244-251.

15. Tsui A, Kuh D, Richards M, Davis D: Delirium symptoms are associated with decline in cognitive function between ages 53 and 69 years: Findings from a British birth cohort study. Alzheimers Dement 2017.

16. Lundstrom M, Edlund A, Bucht G, Karlsson S, Gustafson Y: Dementia after delirium in patients with femoral neck fractures. Journal of the American Geriatrics Society 2003, 51(7):1002-1006.

17. Rudberg MA, Pompei P, Foreman MD, Ross RE, Cassel CK: The natural history of delirium in older hospitalized patients: a syndrome of heterogeneity. Age and Ageing 1997, 26(3):169-174.

18. Shehabi Y, Riker RR, Bokesch PM, Wisemandle W, Shintani A, Ely W, Grp SS: Delirium duration and mortality in lightly sedated, mechanically ventilated intensive care patients. Critical Care Medicine 2010, 38(12):2311-2318.

19. Morandi A, Rogers BP, Gunther ML, Merkle K, Pandharipande P, Girard TD, Jackson JC, Thompson J, Shintani AK, Geevarghese S et al: The relationship between delirium duration, white matter integrity, and cognitive impairment in intensive care unit survivors as determined by diffusion tensor imaging: the VISIONS prospective cohort magnetic resonance imaging study*. Critical care medicine 2012, 40(7):2182-2189.

20. Yang FM, Marcantonio ER, Inouye SK, Kiely DK, Rudolph JL, Fearing MA, Jones RN: Phenomenological Subtypes of Delirium in Older Persons: Patterns, Prevalence, and Prognosis. Psychosomatics 2009, 50(3):248-254.

21. Jackson TA, Wilson D, Richardson S, Lord JM: Predicting outcome in older hospital patients with delirium: a systematic literature review. Int J Geriatr Psychiatry 2016, 31(4):392-399.

22. Mathews SB, Arnold SE, Epperson CN: Hospitalization and cognitive decline: Can the nature of the relationship be deciphered? The American journal of geriatric psychiatry: official journal of the American Association for Geriatric Psychiatry 2014, 22(5):465-480.

23. Ehlenbach WJ, Hough CL, Crane PK, Haneuse SJ, Carson SS, Curtis JR, Larson EB: Association between acute care and critical illness hospitalization and cognitive function in older adults. JAMA: the journal of the American Medical Association 2010, 303(8):763-770.

24. Wilson RS, Hebert LE, Scherr PA, Dong X, Leurgens SE, Evans DA: Cognitive decline after hospitalization in a community population of older persons. Neurology 2012, 78(13):950-956.

25. Matthews FE, Arthur A, Barnes LE, Bond J, Jagger C, Robinson L, Brayne C: A two-decade comparison of prevalence of dementia in individuals aged 65 years and older from three geographical areas of England: results of the Cognitive Function and Ageing Study I and II. Lancet 2013.

26. Sweet L, Adamis D, Meagher DJ, Davis D, Currow DC, Bush SH, Barnes C, Hartwick M, Agar M, Simon J et al: Ethical challenges and solutions regarding delirium studies in palliative care. Journal of Pain and Symptom Management 2014, 48(2).

27. Rhodius-Meester H, van Campen J, Fung W, Meagher D, van Munster B, de Jonghe J: Development and validation of the Informant Assessment of Geriatric Delirium Scale (I-AGeD). Recognition of delirium in geriatric patients. European Geriatric Medicine 2013, 4(2):73-77.

28. Buckwalter JG, Crooks VC, Petitti DB: A preliminary psychometric analysis of a computer-assisted administration of the Telephone Interview of Cognitive Status-modified. J. Clin Exp Neuropsychol 2002, 24(2):168-175.

29. Cook SE, Marsiske M, McCoy KJ: The use of the Modified Telephone Interview for Cognitive Status (TICS-M) in the detection of amnestic mild cognitive impairment. J Geriatr Psychiatry Neurol 2009, 22(2):103-109.

30. Herr M, Ankri J: A critical review of the use of telephone tests to identify cognitive impairment in epidemiology and clinical research. J Telemed Telecare 2013, 19(1):45-54.

31. Plassman BL, Newman TT, Welsh KA, Helms M, Breitner JCS: Properties of the Telephone Interview for Cogntive Status - application in epidemiologic and longitudinal studies. Neuropsychiatry Neuropsychol Behav Neurol 1994, 7(3):235-241.

32. Fleming J, Brayne C: Inability to get up after falling, subsequent time on 4floor, and summoning help: prospective cohort study in people over 90. Bmj 2008, 337:a2227.

33. Gill TM: Disentangling the disabling process: insights from the precipitating events project. Gerontologist 2014, 54(4):533-549.

34. Tieges Z, Stiobhairt A, Scott K, Suchorab K, Weir A, Parks S, Shenkin S, MacLullich A: Development of a smartphone application for the objective detection of attentional deficits in delirium. Int Psychogeriatr 2015, 27(8):1251-1262.

35. Kuhn E, Du X, McGrath K, Coveney S, O'Regan N, Richardson S, Teodorczuk A, Allan L, Wilson D, Inouye SK et al: Validation of a consensus method for identifying delirium from hospital records. PLOS One 2014, 9(11):e111823.

36. Eeles EM, Hubbard RE, White SV, O'Mahony MS, Sawa GM, Bayer AJ: Hospital use, institutionalisation and mortality associated with delirium. Age Ageing 2010, 39(4):470-475.

37. Therneau TM, Grambsch, P.M.: Modeling survival data: extending the Cox model. New York: Springer Verlag; 2000.

38. Henderson R, Diggle P, Dobson A: Joint modelling of longitudinal measurements and event time data. Biostatistics 2000, 1(4):465-480.

39. Skelly D, Griffin EW, Murray C, Harney S, O'Boyle C, Hennessy E, Rawlins N, Bannerman D, Cunningham C: Acute transient cognitive dysfunction and acute brain injury induced by systemic inflammation occur by dissociable IL-1-dependent mechanisms. bioRxiv 2017.

40. Davis DH, Skelly DT, Murray C, Hennessy E, Bowen J, Norton S, Brayne C, Rahkonen T, Sulkava R, Sanderson DJ et al: Worsening cognitive impairment and neurodegenerative pathology progressively increase risk for delirium. The American journal of geriatric psychiatry: official journal of the American Association for Geriatric Psychiatry 2015, 23(4):403-415.

41. Richardson SJ, Davis DHJ, Stephan B, Robinson L, Brayne C, Barnes L, Parker S, Allan LM: Protocol for the Delirium and Cognitive Impact in Dementia (DECIDE) study: A nested prospective longitudinal cohort study. BMC Geriatr 2017, 17(1):98.

42. Kuh D, Pierce M, Adams J, Deanfield J, Ekelund U, Friberg P, Ghosh AK, Harwood N, Hughes A, Macfarlane PW et al: Cohort profile: updating the cohort profile for the MRC National Survey of Health and Development: a new clinic-based data collection for ageing research. International Journal of Epidemiology 2011, 40(1):e1-9.

43. Kuh D, Wong A, Shah I, Moore A, Popham M, Curran P, Davis D, Sharma N, Richards M, Stafford M et al: The MRC National Survey of Health and Development reaches age 70: maintaining participation at older ages in a birth cohort study. Eur J Epidemiol 2016.

44. Fong TG, Davis D, Growdon ME, Albuquerque A, Inouye SK: The interface between delirium and dementia in elderly adults. Lancet Neurol 2015, 14(8):823-832.

45. Richardson S, Teodorczuk A, Bellelli G, Davis DH, Neufeld KJ, Kamholz BA, Trabucchi M, MacLullich AM, Morandi A: Delirium superimposed on dementia: a survey of delirium specialists shows a lack of consensus in clinical practice and research studies. Int Psychogeriatr 2016, 28(5):853-861.

46. Morandi A, Davis D, Bellelli G, Arora RC, Caplan GA, Kamholz B, Kolanowski A, Fick DM, Kreisel S, MacLullich A et al: The Diagnosis of Delirium Superimposed on Dementia: An Emerging Challenge. J Am Med Dir Assoc 2017, 18(1):12-18.

47. Matthews FE, Stephan BC, Robinson L, Jagger C, Barnes LE, Arthur A, Brayne C, Cognitive F, Ageing Studies C: A two decade dementia incidence comparison from the Cognitive Function and Ageing Studies I and II. Nat Commun 2016, 7:11398.

